# MEP-independent silent periods in hand muscles elicited by Transcranial Magnetic Stimulation of the ventral premotor cortex: a non-invasive tool to explore premotor negative motor areas

**DOI:** 10.1101/2025.10.23.684178

**Authors:** PierPaolo Berti, Guido Barchiesi, Mara Turri, Roberta Volpini, Giancarlo Caschera, Roberta Spano, Fabio Mariotti, Iolanda Galdi, Paolo Cipriano Cecchi, Andreas Schwarz, Francesco Sala, Christoph Griessenauer, Luigi Cattaneo

## Abstract

**Objective:** We investigate the possibility to disrupt motor activity via premotor and parietal cortex stimulation, by inducing cortical silent periods (cSPs) in the voluntarily activated upper limb.

**Methods:** We analyzed data from 17 subjects with normal brain function, using navigated TMS (nTMS) on individual MR anatomies. We applied single-pulse biphasic stimulation at 120% of resting motor threshold (rMT) in blocks of 30 stimulations on each spot of a 10–16 point grid covering the inferior parietal and frontal lobes in the dominant hemisphere while participants performed voluntary submaximal contraction. Electromyography (EMG) was recorded bilaterally from intrinsic hand muscles.

**Results:** We observed cSPs not preceded by a MEP in the contralateral hand in 16/17 participants. The maximum overlap, of individual areas where such MEP-independent cSPs could be evoked, corresponded to the ventral precentral gyrus (MNI coordinates: [x=-57, y=7, z=33]). In a subset of stimulus sites, MEP-independent cSPs were bilateral, with contralateral predominance. Canonical short-latency MEPs were observed in all patients, with maximum overlap over the primary motor cortex. We also observed rare contralateral long-latency (> 23 ms) MEP-like responses from peri-Rolandic TMS.

**Conclusions:** MEP-independent cSPs are systematically elicitable in healthy participants They likely reflect interference with premotor representations of ongoing movements. They offer a novel possibility to investigate higher-order motor functions in the experimental and clinical neurosciences.

## 1. Introduction

### 1.1 Negative motor responses (NMRs) and silent periods (SPs)

Negative motor responses (NMRs) are electrophysiological and behavioral phenomena defined as the transient, complete or partial, inhibition of voluntary movement without loss of consciousness (Filevich et al., 2012; Lüders et al., 1992) that are elicited by stimulation of the nervous system. The present work focuses on NMRs evoked by cortical stimulation, although NMRs can be elicited by stimulation of any portion of the central and peripheral nervous system. Cortical NMRs are commonly elicited by means of invasive, direct electrical stimulation (DES) of the cortex during brain surgery or intracranial stimulations. DES-induced negative motor responses have been described since the origins of intraoperative cortical stimulation (Penfield and Boldrey, 1937), but only in recent years has the scientific community become acutely aware of the relevance of NMRs in relation to motor control and cognition (Fornia et al., 2020). NMRs are observed during the voluntary contraction of a muscle and therefore require the subject to be awake and cooperative. The simplest way to observe an NMR is by having the participant perform an isometric contraction. NMRs are documented behaviorally as a drop in active posture when held against gravity (Rubboli and Tassinari, 2006), or instrumentally as a pause in the ongoing EMG pattern. Such pauses in voluntary EMG are referred to as Silent Periods (SPs). SPs are a commonly observed phenomenon in clinical neurophysiology and are the expression of different possible mechanisms. A true SP can occur as a consequence of pre- or post-synaptic inhibition of neurons along motor pathways; however, pseudo-SPs can occur as a consequence of non-inhibitory mechanisms, such as synchronization of motor unit discharge or collision with antidromic stimuli (Floeter, 2003). Given these premises, non-invasive, preoperative mapping of negative motor areas (NMAs) with TMS holds strong potential for informing surgical planning. However, TMS is not as efficient as DES in eliciting pure NMRs. The apparent dissociation between intraoperative stimulation and TMS is likely due to the different stimulation protocols used in the two techniques. Intraoperative mapping usually employs 60 Hz bipolar direct electrical stimulation, delivered in variable trains of 1–5 seconds to identify functional sites (Fornia et al., 2022; Rech et al., 2019). Such stimulation parameters are well beyond the established safety limits for TMS. Indeed, single TMS pulses have rarely been reported to elicit pure NMRs. The functional interpretation of NMAs is still debated. While some authors claim that NMRs are a simple epiphenomenon of electrical stimulation over nodes of the cortical motor system that produce active movements (Mikuni et al., 2006), others claim that they represent a dedicated system for action inhibition (Filevich et al., 2012; Swann et al., 2012). However, the fact that NMAs occur mostly symmetrically in both hemispheres and their widespread distribution in the peri-Rolandic regions (Borggraefe et al., 2016) make it unlikely that they exclusively represent the expression of an action inhibition network. Finally, intraoperatively defined NMAs mostly represent purely inhibitory phenomena, as they are not preceded by positive, excitatory responses, and NMRs tend to have a lower threshold than excitatory responses (Ikeda et al., 2000; Mikuni et al., 2006; Rech et al., 2019; Uematsu et al., 1992). NMAs therefore represent cortical nodes in the action system that produce a pure suppression of motor output when stimulated during an active voluntary task.

From a clinical perspective, NMAs are thought to be indicative of higher-order motor functions such as movement initiation, bimanual coordination, motor awareness, hand selection, and praxis (Fornia et al., 2020; Tariciotti et al., 2024). Lesions in NMAs can produce motor deficits with features distinct from those caused by lesions in the primary motor cortex (Fornia et al., 2022; Giampiccolo et al., 2021; Rossi et al., 2019).

### 1.2 Exploration of Negative motor areas (NMAs) with non-invasive brain stimulation

A TMS-evoked SP occurs following a motor-evoked potential (MEP) after stimulation of the primary motor cortex (M1) (Kofler et al., 2021). Such an MEP-dependent cortical SP (cSP) occurs contralaterally to the stimulated hemisphere but is also observed ipsilaterally to stimulation (ipsilateral SP), where it is the result of transcallosal connections to the contralateral M1 (Zazio et al., 2022). There is general consensus that MEP-dependent cSPs are generated, at least their later components, by cortical inhibitory mechanisms that are different from those producing an MEP (Chen et al., 1999; Davey et al., 1994; Kallioniemi et al., 2014; Kofler et al., 2021; Stetkarova and Kofler, 2013). Some authors have argued that cortical SPs are maximally elicited from regions that are non-overlapping with the peak cortical regions for MEP elicitation (Pitkänen et al., 2015), and one report identified an anterior region on the scalp from which MEP-independent cSPs could be elicited (Wassermann et al., 1993). To summarize the current evidence: a) TMS systematically evokes cSPs in association with MEPs; b) there is general consensus that at least the later components of the cSP are produced by mechanisms independent of those producing the MEP; c) sparse reports indicate that cSPs can be evoked by TMS in the absence of MEPs. It still remains an open question whether cSPs can be obtained by TMS in the complete absence of an MEP by stimulating NMAs in extra-M1 cortical regions, in analogy with the well-known NMRs from invasive stimulation.

### 1.3 Aim of the study

In the present study, we aimed to explore NMAs with TMS coupled with state-of-the-art neuronavigation of individual brain anatomies. We focused specifically on the ventral portion of the frontal and parietal lobes, where we investigated the possible existence of extra-M1, MEP-independent cortical SPs. Importantly, we investigated every participant with a dense grid of TMS targets, covering the cortical regions of interest evenly and with a sufficient spatial sampling rate (Cattaneo, 2018). We evaluated electromyographic changes in hand muscles of healthy participants following suprathreshold single-pulse nTMS applied along the precentral and postcentral gyri, premotor cortex, pars opercularis of the inferior frontal gyrus (IFG), and posterior parietal cortex (inferior parietal lobule and intraparietal sulcus). We consistently observed MEP-independent cSPs in the vast majority of subjects, with maximum overlap in the ventral frontal regions. We hypothesize that developing a feasible, reproducible, and non-invasive mapping technique for NMAs with nTMS also has clinical potential, possibly to enhance preoperative planning, thereby informing treatment decisions and improving patient counselling

## 2. Methods

### 2.1 Participants

We analyzed data from 17 participants (8 male, 9 female; mean age 60), recruited based on the following exclusion criteria: age < 18 years; presence of brain lesions; personal or family history of epilepsy; treatment with neuroactive drugs; and metallic implants in the head and neck. The study was carried out at the neurosurgical center in Bolzano (South Tyrol Health Trust, Italy). All participants provided informed consent and were screened for potential contraindications to TMS, following established safety guidelines (Rossi et al., 2021). The study was approved by the local ethics committee (protocol N. 4/2024 bis) and conducted in accordance with the revised Declaration of Helsinki (World Medical Association, 2013). The participants in the current report were recruited as a neurologically intact control group as part of a broader study protocol, the aim of which is beyond the scope of the present work.

### 2.2 MRI

All patients underwent preoperative 3-Tesla MRI scans, which included the acquisition of T1-weighted 3D MPR, T2-weighted, and FLAIR (echo train length: 48, TE: 83 ms, TR: 6,800 ms, slice thickness: 3 mm, 20 diffusion directions, b-value = 1000). The T1-weighted sequences (echo train length: 1, TE: 2.67 ms, TR: 2,000 ms, matrix resolution: 256 × 246, slice thickness: 1 mm) were typically utilized as the anatomical reference scan. If a T1-weighted image was not available or not compatible with the TMS machine, T2-weighted fast spin-echo sequences (TR/TE: 5200 ms / 100 ms), or T2-weighted inversion recovery fast spin-echo sequences (TR/TE/TI: 6000 ms / 150 ms / 2000 ms) were employed. The raw DICOM images were used to reconstruct a render of the cortical surface using the mriCROn software (Rorden and Brett, 2000). A grid of 12–15 target points was drawn to cover the ventral portion of the frontal and parietal lobes, including the caudal inferior frontal gyrus (IFG), the ventral precentral gyrus (preCG), the ventral postcentral gyrus (postCG), and the inferior parietal lobule (IPL), as shown in **Figure 1**. In six participants, we also applied control stimuli to the occipital lobe and to the midline corresponding to the supplementary motor region, to control for non-specific effects of TMS (i.e., acoustic noise, trigeminal stimulation, mechanical stimulation of the skull).

**Figure 1:**
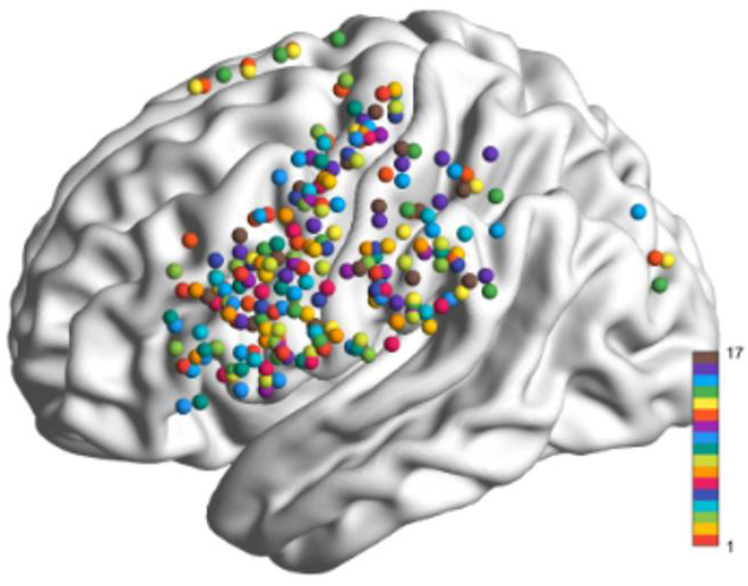
MNI152 standard brain with superimposed TMS spots, represented as 4 mm dimeter spheres-Left: Cumulative grid showing all stimulation spots from the 17 participants (color scale indicates the participant’s number).

### 2.3 nTMS mapping

Single-pulse biphasic TMS was delivered through a figure-of-eight coil (70 mm diameter of external windings) using the Nextim NBS System 5 (Nexstim Oy, Helsinki, Finland). EMG recordings were performed bilaterally from the first dorsal interosseous (1DI) and the Abductor Pollicis Brevis (APB) muscles by means of surface electrodes in a bipolar belly-tendon montage. The raw signal was amplified 1000-fold and digitized at 10 kHz. For the purpose of coregistration between the patient’s scalp and MRI data, anatomical landmarks included the crus of the helix on both sides, the nasion, and nine additional scalp points identified by the nTMS software. The maximum stereotactic error tolerated by the system was 2 mm; stimulation was halted if this threshold was exceeded. The mapping procedure was preceded by the identification of the motor hotspot and assessment of the resting motor threshold (rMT). TMS mapping was performed exclusively on the dominant hemisphere as determined by the Edinburgh Handedness Inventory (Oldfield, 1971). The mapping procedure consisted of stimulating each point, in a random order, with 30 stimuli per spot. The stimulation intensity was suprathreshold and set at 110% of the rMT obtained in the 1DI muscle. To minimize preconditioning effects, both the task interval and the timing of TMS pulses were varied across trials. The behavioral task consisted of an isometric contraction of the hand around a small cylindrical object. Single TMS pulses were delivered manually by the operator whenever a stable contraction was obtained, based on audio and visual inspection of the online surface EMG traces.

### 2.4 Data pre-processing and analysis

The EMG traces were analyzed as event-related epochs, centered on the TMS pulse. The following pre-processing steps were applied: the raw signal was individually inspected to remove artifacts (0% discard rate). The recordings consisted of a single continuous trace for each spot. The data were preprocessed with a custom-written MATLAB R2022b (The MathWorks Inc.; 2024) script and using FieldTrip (Oostenveld et al., 2011). The recording was downsampled to 2 kHz, band-pass filtered between 10 Hz and 1 kHz, and rectified. The rectified trace was smoothed with a 20-sample sliding window.

The analysis started with the segmentation of the recordings into epochs spanning 200 ms before and 200 ms after each stimulus. Single rectified traces were then averaged within each stimulation spot. In this way, we obtained a single rectified averaged trace for each muscle at each TMS spot. Finally, the recordings from 1DI and APB were averaged to obtain a single representative trace of the distal hand muscles. The occurrence of significant post-TMS events in the averaged traces was determined by comparing post-TMS segments with the pre-TMS baseline. To do so, we constructed an “ error box” based on the 200 ms pre-TMS interval: we calculated the mean value of pre-TMS EMG ± 1.96 standard deviations. Any deviation in the post-TMS EMG exceeding the upper and lower bounds of the error box was considered a significant effect of TMS, provided that it lasted for 10 consecutive milliseconds (**Figures 2-5** show examples). Any event exceeding 100 ms in latency was disregarded to exclude contamination by voluntary responses to TMS-evoked sensations. Importantly, only the first post-TMS events were considered. For example, in the case of an MEP followed by an SP, only the primary positive response was considered. Conversely, in the case of a primary inhibition followed by a rebound in EMG activity, as is usually the case for SPs (Floeter, 2003; Türker and Powers, 2005), we considered only the primary inhibitory response. The data were analyzed with a custom-written MATLAB R2022b (The MathWorks Inc.; 2024) code. The code identified significant deviations outside the ±1.96 SD error box and reported the latency of the initial crossing of the limit and the area of the curve outside the error box.

**Figure 2:**
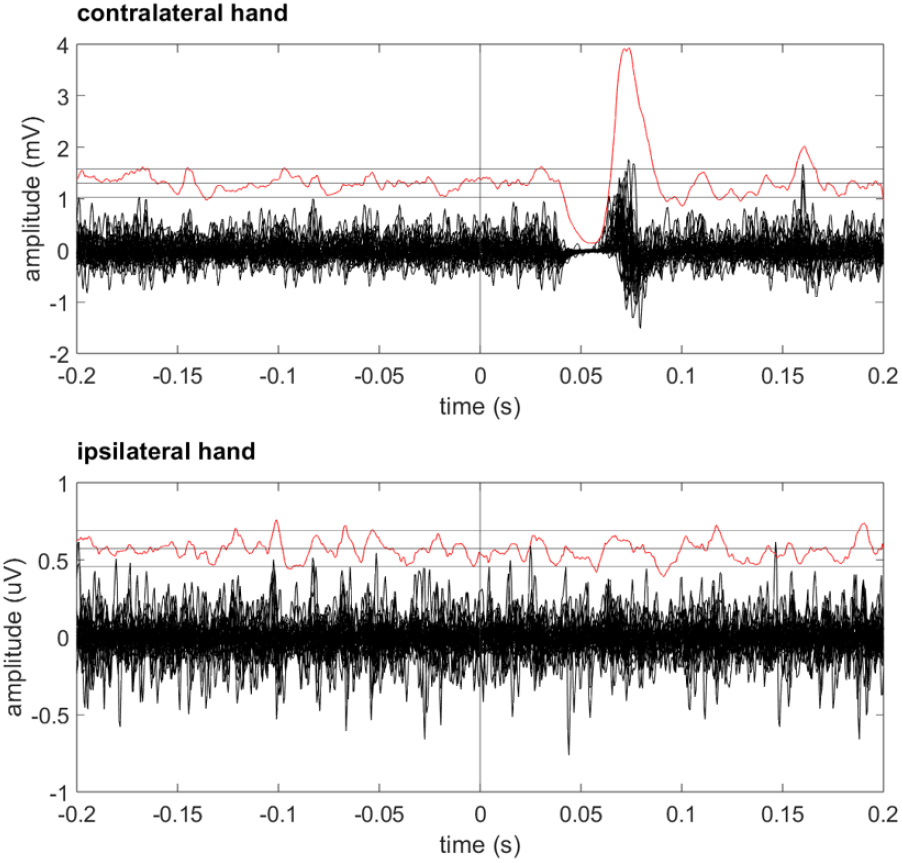
Representative image of a complete cSP from a single subject. (Subjj #13, spot #2). The black lines indicate the recordings from the APB and 1DI muscles superimposed and aligned to the time of TMS (t=0). The red lines represent the rectified, smoothed and averaged compound EMG. The horizontal black lines indicate the “ error box” for the rectified trace (baseline mean +/-1.96*baseline SD). Note that for illustration purposes the rectified trace and corresponding error boxes have been rescaled 5x for appropriate comparison with the raw traces.

### 2.5 Topography of the post-TMS events

All individual brain scans were preprocessed using SPM12 (https://www.fil.ion.ucl.ac.uk/spm/software/spm12/) with the following steps: 1) segmentation and transformation into Montreal Neurological Institute (MNI) standard space to obtain 1 mm voxel MNI-normalized images. The coordinates of the target points, generated as an image in the subject’s native MRI space by the Nexstim neuronavigation system, were transformed into MNI space using the individually obtained transformation matrix. Note that in one left-handed participant, the right hemisphere was stimulated, and the relative coordinates were flipped to the left hemisphere for consistency in the analysis. The final results of the mapping procedure for each individual stimulation spot were, therefore, the set of x, y, and z coordinates in MNI space, the polarity of the effect (inhibitory, null, or excitatory), and the latency and area of the inhibitory responses (SPs). For illustration purposes, the cortical targets were visualized with the BrainNet Viewer (http://www.nitrc.org/projects/bnv/) (Xia et al., 2013). The cumulative grid of all TMS spots is shown in **Figure 1**.

To produce population-level data, we could not use the single spots because TMS targets from a single subject do not represent independent data points. This part of the analysis was carried out with a custom-written Python script, using the Nilearn (Machine learning for NeuroImaging in Python) toolbox (Abraham et al., 2014). To obtain quantitative spatial information at the group level, we first analyzed each subject by extracting the spots that produced a given type of response (e.g., contralateral MEP-independent cSP). We projected these coordinates onto the surface of the pial-left surface template from FreeSurfer and interpolated an area around those spots with a 10 mm margin from each coordinate. All individual surfaces were then superimposed to produce a heatmap of subject overlap, and a local maximum was identified. Finally, the topography of local maxima for each post-TMS effect was compared with the parcellation of the premotor and parietal cortices of the Julich brain atlas (Amunts et al., 2020).

### 2.6 Control experiments

We performed two control conditions. The first was to exclude the possibility that the effects could be due to non-specific TMS features such as noise, electrical stimulation of the scalp, or mechanical head displacement. We therefore applied TMS to targets over the midline of the frontal region and over the occipital cortex. In another subset of 8 participants, we constructed a recruitment curve of the MEP and of the MEP-associated cSP, testing several stimulus intensities over M1, ranging from 70% to 130% of the resting motor threshold. This control was performed to exclude the possibility that subthreshold stimulation of M1 could elicit a cSP in the absence of an MEP.

## 3. Results

None of the participants reported any immediate or delayed side effects from the stimulation. We observed several patterns of EMG response to TMS in the voluntarily activated muscles. In the hand contralateral to the stimulated hemisphere, we found: A) short-latency MEPs, compatible with canonical fast corticospinal conduction (latency < 24 ms); B) MEP-independent cSPs, i.e., pauses in voluntary EMG not preceded by excitatory phenomena; C) long-latency (>24 ms) excitatory phenomena; and D) no effect. In the hand ipsilateral to the stimulated hemisphere, we observed three possible phenomena: A) cSPs associated with contralateral MEPs; B) cSPs not associated with contralateral MEPs; or C) no effect. An illustration of paradigmatic cases for each condition is provided in **Figures 2-5**.

### 3.1 Contralateral MEP-independent cSP

We found evidence of MEP-independent cSPs in at least one spot of the grid in 94% of participants (16/17 subjects; subj #5 did not show any). The number of grid spots from which a cSP could be elicited varied from 1 to 9, indicating considerable individual variability in the spatial extent of the NMRs. The peak overlap corresponded to the anterior portion of the precentral gyrus, at the junction with the inferior frontal sulcus at the MNI coordinates of [x=-57, y=7, z=33], but with considerable overlap along the mid-portion of the precentral gyrus (**Figure 6A**). The mean onset latency of the contralateral MEP-independent cSPs was 36.7 ms (SD = 8.0 ms). The distribution of the onset latency was unimodal, as illustrated in **Figure 7A**. The duration of the MEP-independent cSP was variable, ranging from 10 to 42 ms. The mean duration was 18.1 ms (SD = 6.8 ms). The decrease in voluntary activity was complete in only a minority of cases, while in the majority, it was an incomplete suppression of EMG (cf. **Figures 2 and 3**). The maximum overlap spot co-localized with the 6v3 parcellation of the Julich atlas (Amunts et al., 2020).

**Figure 3:**
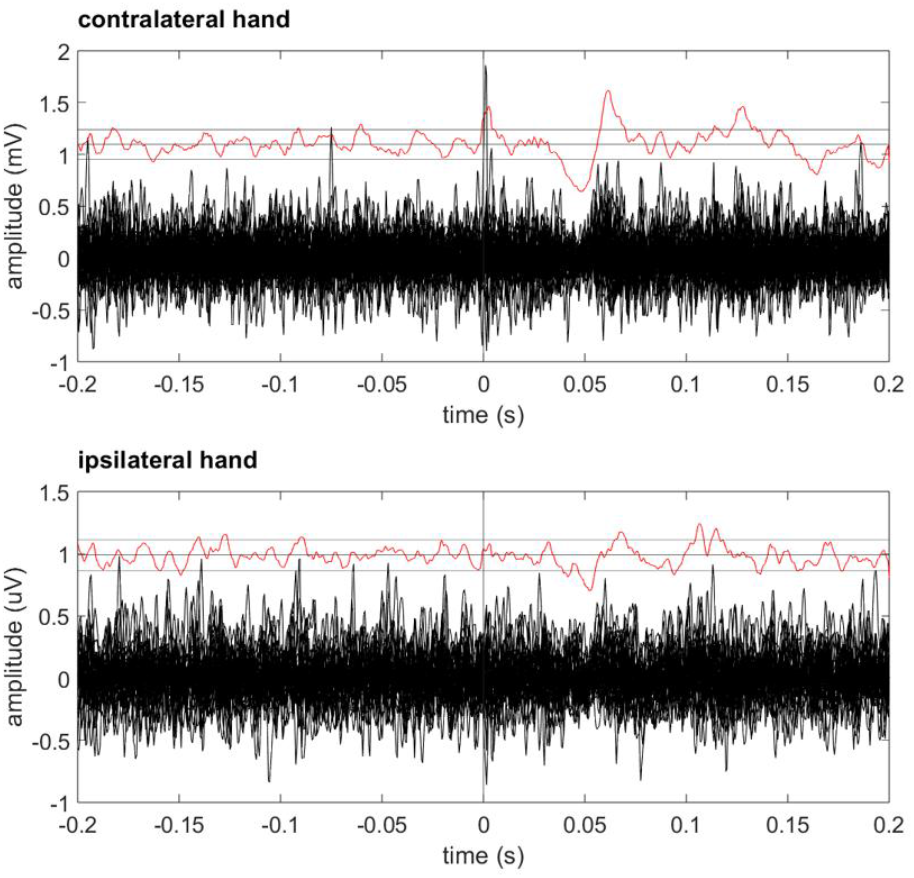
Representative image of a partial cSP from a single subject. (Subj #2, spot #2). Conventions as in Fig. 2

### 3.2 Contralateral MEPs

As expected, all subjects showed excitatory responses compatible with MEPs elicited from the Rolandic region by suprathreshold TMS. The latency of the MEPs showed a bimodal distribution, with an early peak around 17 ms and a late peak around 25 ms (**Figure 7B**). We defined the cut-off value between early and late responses as 23 ms, based on literature descriptions of early response latency (Groppa et al., 2012) and on visual inspection of the present data. **Figure 4** shows a typical early-onset MEP, and **Figure 5** shows a typical long-latency MEP. MEPs belonging to the early peak corresponded to MEPs elicited from M1, were evident in all participants, and showed maximum overlap on the hand-knob of the precentral gyrus at the coordinates [x=-37, y=-18, z=69] (**Figure 6B**). MEPs with latencies in the late peak were much less frequent (present in 7/17 participants) and did not show a consistent spatial location in the peri-Rolandic region, being distributed between the parietal, frontal opercular, and premotor regions. Their population map failed to show a consistent spatial location, with an insignificant peak overlap of only two subjects out of seven at the coordinates [x=-59, y=-32, z=42].

**figure 4:**
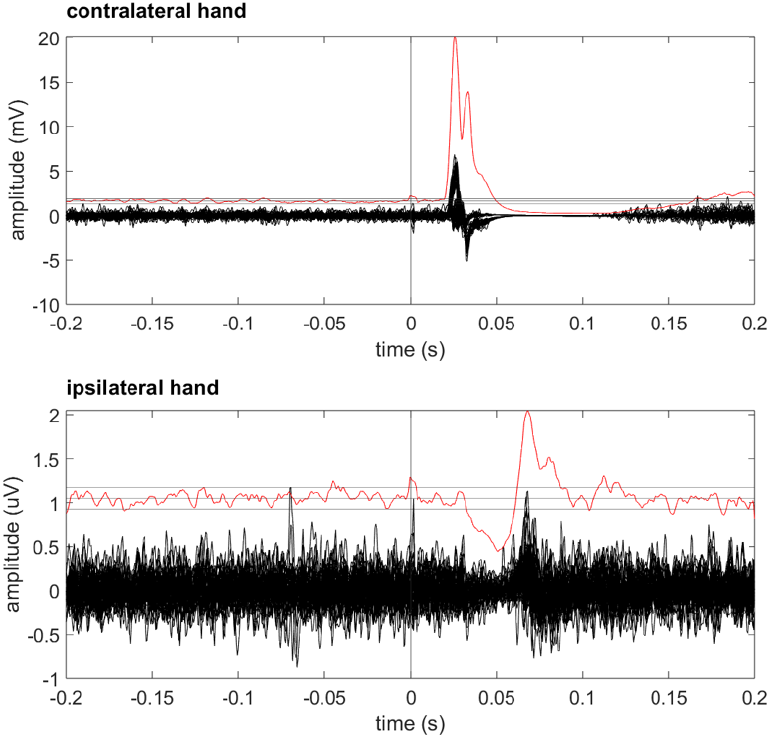
Representative Image of a short-latency MFP from a single subject. (Subj #2, spot #l). Conventions as in Fig. 2

**Figure 5:**
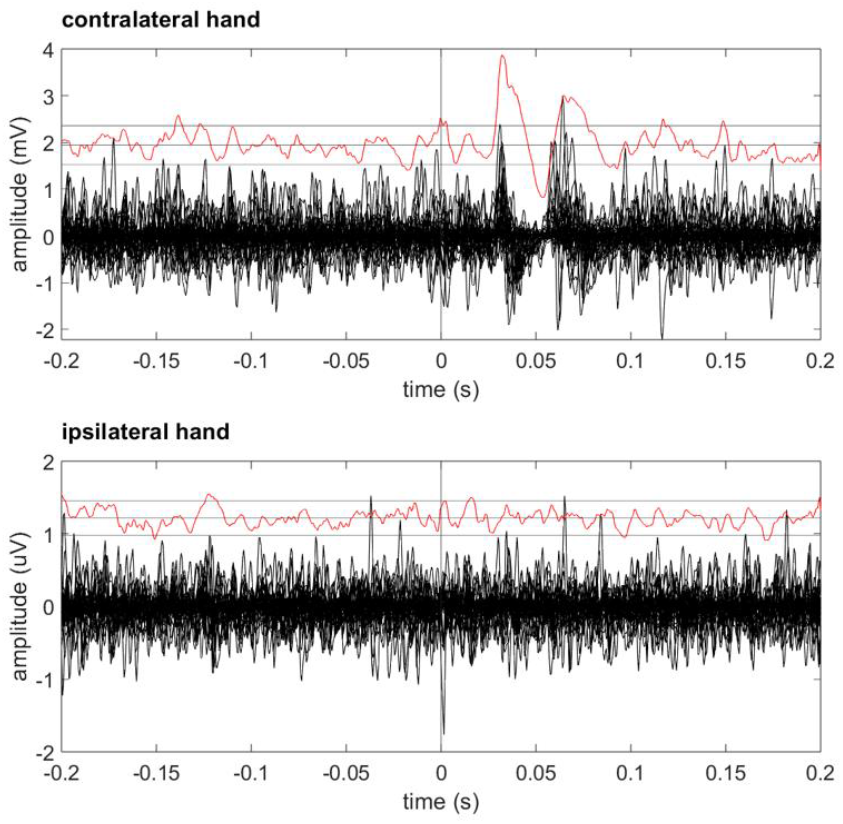
Representative image of a long-latency MEP from a single subject. (Subj #15, spot #5). The MEP shown here has an onset latency of 29 ms. Conventions as in Fig. 2

**Figure 6:**
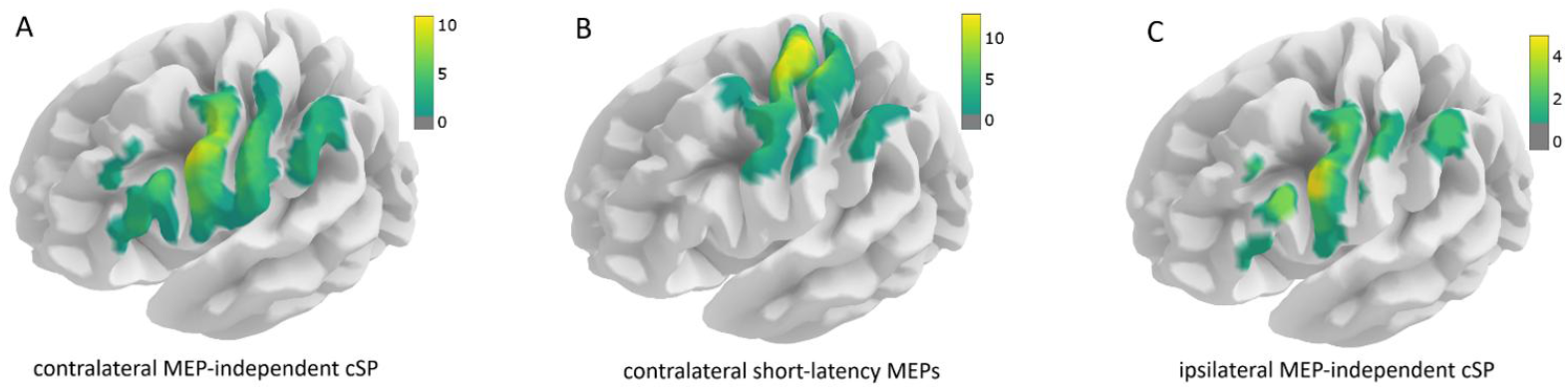
overlay maps of the number of participants showing significant effects of stimulation in a given region. Please note that scales differ between sub-plots and therefore direct comparison of colormaps should not be made.

### 3.3 Ipsilateral silent periods

We observed the systematic presence of the well-known ipsilateral silent periods (Bortoletto et al., 2021; Ferbert et al., 1992) in association with contralateral short-latency MEPs. The description of these is beyond the scope of the present work. Unexpectedly, we also observed the presence of ipsilateral cSPs independent of contralateral MEPs. These were never complete SPs and were observed only in association with contralateral MEP-independent cSPs; they were systematically smaller and were present in only 9/17 participants. **Figure 3** illustrates the simultaneous presence of two incomplete inhibitions of EMG activity, with a prevalence on the side contralateral to TMS. The mean latency of ipsilateral MEP-independent cSPs was 45.8 ms (SD = 12.0 ms), and the mean duration was 17.8 ms (SD = 8.7 ms). In no case did we find standalone ipsilateral cSPs without any contralateral phenomenon. The overlap of MEP-independent ipsilateral cSPs was found around the peak coordinates of [x=-57, y=8, z=31], which nearly overlapped with those for the MEP-independent contralateral cSP. It should be noted that we never observed ipsilateral excitatory responses.

### 3.4 Control experiments

Regarding the first control condition, in none of the frontal and occipital control sites did we find TMS-evoked activity, neither excitatory nor inhibitory. The second control experiment showed that in none of the subjects (n=8), TMS over M1 elicited a cSP in the absence of a preceding MEP (**Figure 8**).

**Figure 7:**
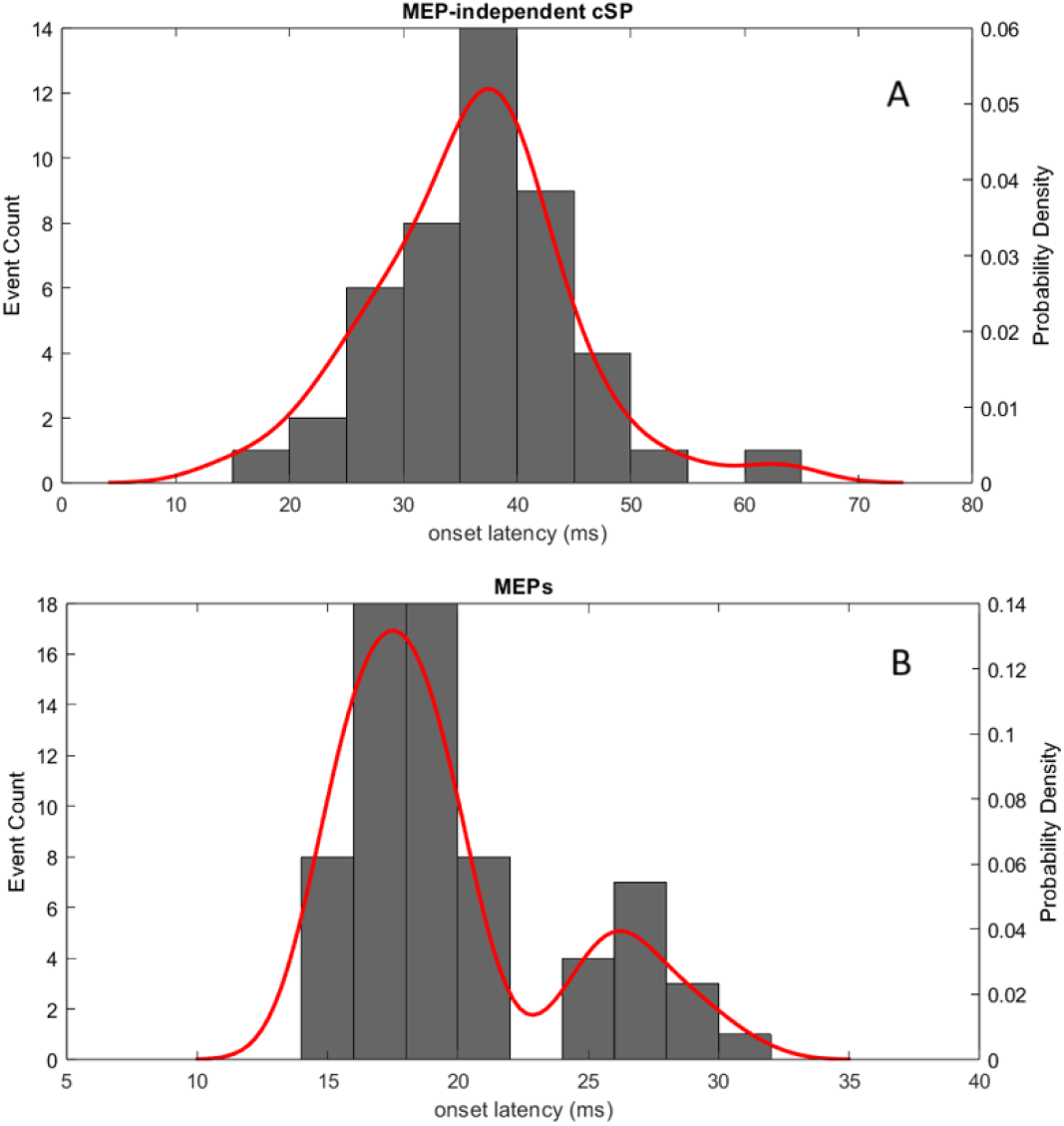
distribution of the number of TMS-evoked responses according to latency of onset. The grey columns indicate raw event counts and the red line indicates the kernel density estimation of event counts. The top panel refers to contralateral MEP-independent cSPs. The bottom panel refers to contralateral excitatory responses, clearly indicating a biphasic distribution of onset latencies. Note that the time scale (x axis) differs between the two graphs./

**Figure 8:**
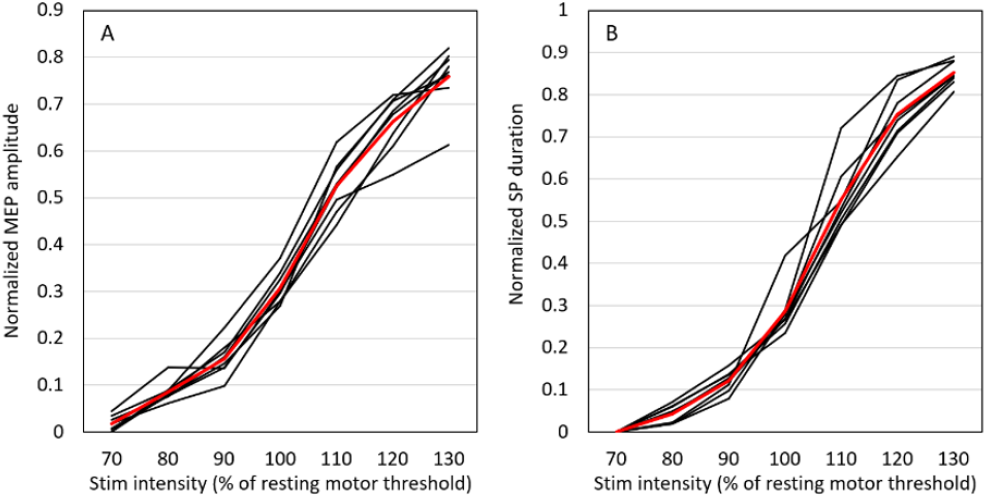
Results of control experiment 2: recruitment curve of the MEP amplitude and of the MEP-dependent cSP in 8 participants.

## 4. Discussion

### 4.1 MEP-independent contralateral cSPs are genuine evoked responses to cortical stimulation

The present study provides novel evidence that cSPs can be consistently evoked from activated hand muscles after stimulation of the contralateral ventral premotor regions using single-pulse TMS. In only one participant out of 17 were no significant cSPs elicited. The first relevant question concerns their specificity to cortical stimulation. Indeed, SPs in ongoing voluntary EMG can occur following sensory stimulation (Cattaneo et al., 2007; Floeter, 2003) and therefore could result from the activation of cutaneous afferents from the scalp. However, two lines of evidence in our study support the cortical origin of the cSPs. First, if cSPs were the result of non-specific sensory stimulation (mechanical or acoustical), they would have been consistently recorded from every stimulation site. Instead, by adopting a large stimulation grid, we demonstrated that SPs were elicited only from a limited subset of sites. Additionally, in some participants, we stimulated control regions far from the parieto-frontal cortex (i.e., the midline and the lateral occipital cortices), and these stimulations also showed no significant evoked EMG responses. Second, most responses were either exclusive to or predominant on the side contralateral to stimulation, making it highly unlikely to be a reflex behavior, which would be expected to be either bilateral or ipsilateral to the stimulated site. Finally, voluntary responses were also excluded by the fact that any event-related EMG phenomenon with a latency exceeding 100 ms was excluded from further analysis, thus cutting off any voluntary reaction to the TMS.

### 4.2 MEP-independent contralateral cSPs are spatially dissociated from MEPs and constitute a separate functional entity

We propose that the cSPs described here are a different entity from the MEP-associated SPs that are commonly described following MEPs. We base this statement on anatomical and functional grounds. The hotspot for eliciting MEP-independent cSPs was located in the ventral anterior portion of the precentral gyrus, around the intersection between the precentral sulcus and the inferior frontal sulcus, which was at a 51 mm Euclidean distance from the M1 hotspot for hand MEPs. However, the mere observation that cSPs can be evoked from a region different from that eliciting MEPs is not sufficient to demonstrate true anatomical or functional segregation. This is because the MEP-associated SP might be elicited at a lower threshold than the MEP itself. Thus, the apparent presence of an SP-only region at the periphery of the MEP area could simply reflect current spread and threshold effects rather than distinct functional localization. The control experiment performed on eight participants showed that the MEP-associated SP does not have a lower threshold of elicitation compared to the MEP, and therefore, the premotor MEP-independent cSPs are not likely due to current spread to M1. The spatial localization of the area from which a cSP could be evoked was significantly variable between individuals, ranging from 1 spot to 7 spots per subject. We interpret such variability in terms of stimulus intensity and current spread to the cortices that generate the MEP and the cSP, similarly to how MEPs were obtained in some participants over an extended frontal and parietal region due to current spread to M1 (see, for example, Holmes et al., 2024). On the other hand, the presence of an MEP masks the possible presence of a MEP-independent cSP, and not vice-versa. Therefore, the extent of the area from which a cSP can be elicited depends on current spread to the precentral gyrus (the cSP hotspot) but also, critically, on current spread to M1 (the MEP hotspot).

### 4.3 Functional interpretation

The current data do not allow for an interpretation of the functional role of the neural system that, when activated by TMS, produces the cSP in the ongoing voluntary contraction. The region of the precentral gyrus around the precentral-inferior frontal sulcus junction is located within the ventral premotor cortex (PMv), which is a pivotal node in parieto-frontal circuits controlling goal-directed upper limb movements. The role of the PMv is to translate the multisensory representations of objects and space built up in the parietal cortex into complex motor plans. More specifically, the premotor region at the level of the inferior frontal junction has been implicated in the direct control of hand movements, as shown by several TMS studies that demonstrated its direct connections to M1, carrying relevant information for object-directed actions (Bäumer et al., 2009a; Cattaneo and Barchiesi, 2011; Davare et al., 2009; Maule et al., 2015). The cSP hotspot is also in close proximity to the inferior frontal junction, a region hypothesized to be a critical cortical hub for cognitive control, integrating information from attentional and motor systems. It enables flexible, goal-directed behavior by mediating processes such as task switching, response inhibition, and updating working memory. Its activity is crucial for rapidly adjusting behavior in response to changing environmental demands and internal goals (Brass et al., 2005; Derrfuss et al., 2009).

The precentral cortex could exert its influence on spinal motor neurons to produce the cSP by two mechanisms: either directly via corticospinal projections or indirectly via cortico-cortical projections to M1 or through cortico-basal ganglia-cortical loops. The current data do not allow us to discern between these three possibilities, though the long onset latency of the cSPs and the considerable jitter in their onset time make a direct corticospinal effect unlikely (though do not exclude it). The indirect pathway through M1 is also favored by current models of premotor function in primates that postulate a cortico-cortical pathway to M1 as the main mechanism by which the PMv produces actual movement (Cattaneo et al., 2005; Davare et al., 2009; Shimazu et al., 2004). The current findings are not sufficient to hypothesize a specific functional role for the cSP phenomenon, because of the simplicity of the behavior tested (isometric contraction of the index finger and thumb) and the lack of control tasks. However, we should resist the simplistic interpretation that a negative phenomenon (the EMG inhibition) necessarily indicates an inhibitory function of the underlying neural system. As a matter of fact, TMS evokes non-physiological neural activity that commonly produces “ loss of function” effects, with variable impacts on behavior.

### 4.4 Comparison to previous literature on non-invasive stimulation

Our findings resonate with the early observations of Wassermann and colleagues (1993), who first described silent periods occurring without preceding MEPs. Both studies report short-latency inhibitory phenomena (mean onset ≈ 27 ms in Wassermann et al. vs. ≈ 37 ms in our cohort) with relatively brief durations and often incomplete suppression of EMG activity, occasionally followed by a late excitatory rebound. Importantly, both studies demonstrate that these events are not ubiquitous but arise from selective cortical loci, supporting the notion that inhibitory silent periods and excitatory MEPs reflect distinct physiological substrates. However, several differences distinguish our work from this earlier report. Methodologically, we employed individual MRI-guided neuronavigation in a larger cohort (n = 17), systematically covering the entirety of the ventral parietal, central, and frontal regions with a dense grid of stimulation points. Topographically, our data indicate a specific anatomical hotspot for MEP-independent SPs in the anterior precentral gyrus near the inferior frontal sulcus, which is consistent with the previous findings by Wassermann et al. (1993).

Although direct online stimulation of NMAs with TMS is not efficient in eliciting SPs, alternative techniques such as dual-coil TMS have consistently shown that extra-M1 cortical regions exert an inhibitory effect on corticospinal output (Byblow et al., 2007; Civardi et al., 2001; Koch et al., 2008; Mars et al., 2009; Maule et al., 2015; Parmigiani et al., 2018, 2015). In several dual-coil TMS experiments, conditioning stimuli have been applied to the ventral premotor cortex, in close proximity to the cSP hotspot found in the present data, showing short-latency inhibitory effects on corticospinal excitability to hand muscles at rest (Bäumer et al., 2009b; Cattaneo and Barchiesi, 2011; Davare et al., 2009; de Beukelaar et al., 2016; Maule et al., 2015). Such dual-coil findings are strongly supportive of the indirect origin of the cSPs observed in the present work.

### 4.5 Coherence of current findings with intraoperatively defined NMAs

The spatial location of the MEP-independent cSP shows striking concordance with the distribution of NMRs reported in previous intraoperative mapping studies. NMAs are found in several distinct regions across the medial and lateral surfaces of both hemispheres, with a specific focus on the supplementary motor area (SMA), pre-SMA, the entire precentral gyrus (PreCG), and the caudal inferior frontal gyrus (IFG). Rech et al. (2019) focused on the precentral gyrus during awake surgery in 117 patients and reported behavioral NMAs related to upper limb function within areas 6v, 6d, and 55b of the Glasser atlas parcellation of the human cerebral cortex, a distribution that is highly consistent with our findings. Recent intraoperative works have emphasized that NMRs are not limited to behavioral arrest phenomena but can also be captured through the analysis of electromyographic (EMG) patterns, which may uncover distinct underlying neural substrates (Viganò et al., 2021; Fornia et al., 2020, 2022). Viganò et al. (2021) provided a cortical surface probability-density map of hand NMRs, integrating data from three independent series (Fornia et al., 2020; Simone et al., 2020; Viganò et al., 2019). Their analysis revealed a maximal density of NMRs in the dorsal premotor cortex, particularly near the junction with the inferior frontal gyrus—a distribution highly superimposable on the cluster of MEP-independent SPs described in our cohort. The differentiation between dorsal and ventral premotor contributions, as characterized by Fornia et al. (2020), further enriches this interpretation. Their EMG-based analyses demonstrated that dorsal premotor sites were preferentially associated with recruitment effects and corticomotoneuronal circuit activation, whereas ventral premotor sites were more commonly linked to suppression effects, suggesting a functional specialization for motor execution and sensorimotor integration that is fully compatible with the present results. The robustness of this spatial convergence across studies using distinct methodological approaches (behavioral vs. EMG vs. SP analysis) strongly suggests that the anterior precentral gyrus and adjacent premotor areas constitute a reproducible NMA.

### 4.6 Ipsilateral responses: MEP-dependent and MEP-independent cSPs

Bilateral recordings of hand muscles allowed us to evaluate evoked responses ipsilateral to the stimulated hemisphere. We observed the well-known ipsilateral cSPs (Chen et al., 2003; Zazio et al., 2022), which are considered an expression of transcallosal inhibitory connections between the two primary motor cortices and are necessarily associated with a contralateral MEP (Ferbert et al., 1992). Indeed, we found that most TMS spots that produced an MEP were associated with an ipsilateral cSP. However, we also found ipsilateral MEP-independent cSPs, which were always associated with contralateral MEP-independent cSPs. In no recordings did we find “ standalone” ipsilateral cSPs that were independent of contralateral responses. The topography of the hotspot for ipsilateral MEP-independent cSPs overlapped with that of the contralateral MEP-independent cSP on the ventral precentral gyrus. Taken together, these results show that the MEP-independent response was mainly contralateral but sometimes bilateral, associated with a less consistent ipsilateral inhibitory response. This finding is in agreement with data on NMRs obtained by invasive stimulation, which seem to be more lateralized to the contralateral hand when elicited by stimulation of the precentral gyrus (Borggraefe et al., 2016).

### 4.7 Late-onset contralateral MEPs

Figure 7. clearly illustrates that the contralateral excitatory responses (MEPs) belonged to two different categories according to their onset latency: the short-latency MEPs, compatible with the activation of fast corticospinal axons, were the vast majority of evoked positive responses. However, a second class of MEPs became apparent, with a latency that was incompatible with normal fast corticospinal conduction from M1 (i.e., > 23 ms, as per Groppa et al., 2012). Late MEPs were uncommon events and, probably due to their low frequency, failed to show a distinct overlap hotspot. This finding in the context of the current experiment is serendipitous and requires better definition in further explorations.

## 5. Conclusions

In this study, we describe MEP-independent cSPs evoked by TMS. They consist of a transient inhibition of ongoing voluntary EMG, which occurs mainly contralaterally to the stimulated hemisphere but may also appear as a bilateral, though predominantly contralateral, response. The topographic and neurophysiological characteristics of the cSP indicate that they are a separate entity compared to the cSP commonly observed following an MEP. These findings have potential direct clinical applications in neurosurgical planning. There is increasing attention in the neurosurgical onco-functional balance to higher-order motor functions, and NMAs are currently seen as useful brain regions to evaluate and quantify these functions. The possibility of eliciting MEP-independent cSPs expands the toolbox of non-invasive brain stimulation in the study of the cortical motor system by adding the exploration of NMAs. In this respect, our results bridge a long-standing gap between non-invasive brain stimulation and intraoperative mapping. Taken together, these results open new avenues for the use of TMS in the functional mapping of NMAs, with potential implications for both basic neurophysiology and clinical practice, particularly in the context of preoperative planning for neurosurgical patients.

## Notes

### Competing Interest Statement

The authors have declared no competing interest.

